# Intergenerational effects of maternal and paternal childhood maltreatment on offspring cortical gyrification are sex-specific: the Transmit Radiant Individuality to Offspring (TRIO) study

**DOI:** 10.1101/2025.08.25.671904

**Authors:** Ryo Yamaguchi, Izumi Matsudaira, Yasuyuki Taki

## Abstract

The effects of parental childhood maltreatment (CM) can be transmitted across generations; however, the underlying sex-specific biological mechanisms are poorly understood. To better understand how parental adversity is passed on to offspring, we investigated the impact of maternal and paternal CM on the local gyrification index (LGI) of offspring using parent–offspring trio data. The LGI was selected as a key neurodevelopmental metric because it is thought to reflect the fetal environment and remain relatively stable postnatally, making it less susceptible to later life experiences. Using multiple regression, we analyzed sex-stratified associations between parental CM subtypes and the offspring LGI and tested for mediation by parental psychosocial functioning. The results revealed a distinct parent-of-origin, sex-specific pattern: maternal emotional abuse was associated with a lower LGI in male offspring, whereas paternal physical neglect was associated with a lower LGI in female offspring. These associations were not mediated by the current mental health or marital relationships of the parents. Collectively, these findings provide compelling evidence that the neurodevelopmental consequences of parental CM can be biologically embedded in the next generation even before birth.

## 1. INTRODUCTION

Childhood maltreatment (CM) by caregivers, including abuse and neglect, has lifelong adverse effects on the physical and mental health of victims (1–3). Systematic reviews and meta-analyses have reported robust associations between CM and alterations in brain development (4) (5), behavioral problems (1) and an increased risk of psychiatric disorders (6). Furthermore, females with a history of CM face an increased risk of perinatal complications (7) and elevated parenting stress when they become mothers (8). Despite the stark paucity of research on males compared with females, several adverse effects of CM in males have also been reported (9). Biologically, CM is associated with epigenetic changes in human sperm (10–12). Psychosocially, males with a history of adverse childhood experiences (ACEs) show greater anxiety and depression during their partner’s pregnancy (13) and subsequently experience greater parenting stress postpartum (14). Furthermore, CM is related to negative outcomes in couples, such as lower relationship satisfaction, intimate partner violence, and psychological distress (15). Beyond the impact on the victims themselves, early-life adversity also influences offspring development (16–18). A systematic review and meta-analysis revealed that parental ACEs are associated with offspring outcomes such as internalizing and externalizing symptoms (18). Furthermore, this association was not moderated by sociodemographic factors (parent and offspring age, income, offspring sex, or racial/ethnic minority status) or methodological differences. These findings emphasize the necessity of elucidating the mechanisms of intergenerational transmission (19).

The mechanisms underlying this intergenerational transmission are multifaceted and involve a complex interplay of biological, psychosocial, and environmental factors. As it is an intermediate phenotype (20,21), neuroimaging should be effective for investigating the putative mechanisms of intergenerational transmission. Indeed, a growing body of neuroimaging research, although focused almost exclusively on mothers, has consistently linked maternal CM and ACEs to widespread alterations in offspring brain structure (e.g., gray and white matter volume (GMV)) and function (e.g., functional connectivity), particularly in regions critical for emotion regulation and stress response, such as the amygdala, hippocampus, anterior cingulate cortex, and prefrontal cortex (22–31). Notably, paternal contributions remain largely absent from this literature, with the studies of Karlsson and colleagues representing a rare exception, providing seminal evidence that paternal CM is also associated with alterations in neonatal brain structure (32–34). Taken together, while these studies confirm that parental adversity impacts the developing brain of offspring, several challenges remain in elucidating its underlying biological mechanisms.

A primary gap in the literature is the insufficient attention given to sex differences. As previously discussed, research on the impact of parental adversity on offspring has been overwhelmingly focused on mothers, leaving a significant gap in our understanding of the role of fathers (9,35–37). Furthermore, despite established knowledge of distinct sex differences in brain development (38–40) and reports from animal models suggesting sex-dependent stress sensitivity (41), it remains unclear whether the effects of parental CM differ on the basis of the sex of the offspring (17). Thus, these gaps in the literature underscore the critical importance of investigating effects separately, considering the sex of both the parent and the offspring (9,42,43).

A second key limitation of the literature is its predominant focus on highly plastic brain metrics. Many prior studies have focused on infants and toddlers, a developmental stage of ongoing brain maturation, leaving the long-term consequences of parental adversity in adolescence and adulthood largely unclear (37). Furthermore, commonly targeted metrics, such as the GMV of the amygdala and hippocampus, are known to be highly plastic and susceptible to significant changes from postnatal environmental influences (44,45). It is therefore advantageous to focus on a metric that is less sensitive to these confounding effects of environment and age. Gyrification, the cortical folding process that results in the formation of gyri and sulci to maximize surface area within a limited volume, is essential for higher-order cognitive functions in the human brain (46). The pattern of cortical gyrification is formed during the late second and third trimesters and is thought to remain relatively stable postnatally (47). The pattern of cortical gyrification is distinguished by its high sensitivity to perinatal influences, including genetic and prenatal environmental factors, coupled with its relative stability against postnatal environmental effects. Accordingly, it holds significant promise as a potential biomarker—or endophenotype—for psychiatric disorders such as schizophrenia and major depressive disorder (48–50). Prior studies have shown an association between prenatal factors and gyrification. For example, a study by Mareckova *et al*. (2020) reported an association between maternal prenatal stress and the local gyrification index (LGI) in young adult offspring (51). Furthermore, the LGI has been linked to various perinatal factors, including preterm birth (52), maternal alcohol consumption during pregnancy (53), obstetric complications (54), and childbirth during the COVID-19 pandemic (55). Therefore, the LGI could offer valuable insights into the mechanisms underlying the intergenerational transmission of parental CM effects on offspring brain development.

In the present study, we investigated the association between parental CM and the offspring LGI in parent– offspring trios comprising offspring from adolescence to young adulthood and their biological parents. It has been reported that parental CM affects parental psychological distress and their relationships with their partners (15,37,56). However, we hypothesized that parental CM would be directly associated with the offspring LGI rather than mediated by such parental behavioral factors at the time of data collection because the LGI is a brain structure, which undergoes substantial maturation in early life. To examine sex-specific differences in this association, we conducted analyses separately for each parent–offspring sex combination (father–son, father–daughter, mother–son, and mother–daughter).

## 2. MATERIALS & METHODS

### 2.1. Participants

This research is part of the Transmit Radiant Individuality to Offspring (TRIO) study, an ongoing project (57). This project adhered to the principles enshrined in the Declaration of Helsinki and received approval from the Institutional Review Board of Tohoku University (Approval No. 2022-1-534). Written informed consent was obtained from all participants before the study. For minors (aged <18 years), parental consent was also required. The detailed method for participant recruitment was described in our previous paper (57). This study focuses on a subsample of 152 parent–offspring trios, consisting of offspring aged 15 years and older and their parents.

To control for genetic background, participation was limited to Japanese individuals with no relatives within the third degree of kinship from other ethnicities. Individuals with a history of cerebrovascular disease, brain tumor, intracranial disease, degenerative brain disease, epilepsy, severe heart disease, and brain injury with impaired consciousness were not eligible for inclusion. These conditions were verified at the time of participation, and if any trio member met the exclusion criteria, their participation was excluded.

### 2.2. Parental childhood maltreatment

We quantified each parent’s history of CM by administering the Childhood Trauma Questionnaire (CTQ) (58). The CTQ is a widely used, standardized 28-item self-report measure for retrospectively assessing experiences of childhood abuse and neglect. The CTQ comprises five subscales: emotional abuse (EA), physical abuse (PA), sexual abuse (SA), emotional neglect (EN), and physical neglect (PN). Each subscale consists of five items rated on a 5-point Likert scale, with total scores for each subscale ranging from 5 to 25. For each subscale, a higher score is indicative of a greater severity of CM. We used the Japanese version of the CTQ (CTQ-JNIMH), the reliability and validity of which were established by Nakajima *et al*. (2022) (59). We also assessed the CM experiences of the offspring using the same instrument. This was done to statistically account for the influence of their own maltreatment history, a potential confounder due to the known intergenerational cycle of abuse (60,61).

### 2.3. Parental behaviors

Current levels of non-specific psychological distress of parents were assessed using the Kessler 6 (K6) scale (62). For this study, we administered the Japanese version, the reliability and validity of which have been established (63). The K6 is a 6-item self-report questionnaire that asks respondents to rate the frequency of psychological distress symptoms over the past 30 days. Each item is rated on a 5-point Likert scale (from 0 = “None of the time” to 4 = “All of the time"), yielding a total score ranging from 0 to 24. Higher scores on the K6 indicate greater psychological distress.

Parental attachment style was assessed using the Experiences in Close Relationships–Relationship Structures (ECR-RS) questionnaire (64). For this study, we administered the Japanese version (65) and, as per our research question, utilized only the subscale pertaining to romantic partners. This 9-item subscale yields scores for two dimensions: attachment-related anxiety (ANX) and avoidance (AVO). Each item is rated on a 7-point Likert scale, with higher scores reflecting greater attachment insecurity in each respective dimension.

### 2.4. Image acquisition

All MRI data were collected using a 3-Tesla dStream Achieva scanner (Philips Medical Systems, Best, Netherlands) and a 20-channel head–neck coil. For each participant, T1- and T2-weighted images were acquired.

T1-weighted images (T1WIs) were obtained using a three-dimensional magnetization-prepared rapid gradient echo (MPRAGE) sequence (368 × 368 matrix, repetition time = 11 ms, echo time = 5.1 ms, field of view = 256 × 256 mm, slices = 257 slices, and slice thickness = 0.7 mm), and sagittal T2-weighted images (T2WIs) were obtained with a spin‒echo sequence (368 × 368 matrix, repetition time = 2500 ms, echo time = 3200 ms, field of view = 256 × 256 mm, slices = 250 slices, and slice thickness = 0.7 mm).

Image quality was visually inspected immediately following acquisition, and reimaging was conducted if significant motion artifacts were identified.

### 2.5. Preprocessing of brain images

MRI preprocessing followed our previous study (66). Briefly, AC-PC alignment (67), nonuniform intensity normalization (68), and brain extraction (69) were performed. Surface-based preprocessing was subsequently performed using the FreeSurfer pipeline to compute the LGI at each vertex on the cortical surface (70,71). In the subsequent processing step, cortical parcellation was applied on the basis of the Human Connectome Project Multi-Modal Parcellation version 1.0 (HCP-MMP1) atlas, which consists of 180 anatomically and functionally defined regions per hemisphere (72).

### 2.6. Statistical Analysis

To test the association between parental CM and the offspring LGI, we performed a series of multiple regression analyses. Given that the differential effects of various CM types remain unclear (17), we entered the five CTQ subscale scores for both mothers and fathers as distinct predictor variables in these models. The outcome variable was the LGI for each of the 360 bilateral cortical regions, which was parcellated according to the HCP-MMP1.0 atlas (72). All the models were adjusted for offspring age and their own CTQ total score as covariates. The resulting *p*-values were corrected for multiple comparisons using the Benjamini–Hochberg (BH) method to control the false discovery rate (FDR) (73), with the significance threshold set at a *q*-value < 0.05.

Next, to investigate potential indirect pathways, we conducted a series of multiple regression analyses. We first examined the association between parental CM subtypes and parental psychosocial functioning, which included their psychological distress (K6) and attachment styles (AVO and ANX). We then examined the associations between these parental psychosocial factors and the offspring LGI while adjusting for offspring age. All *p*-values from these analyses were corrected for multiple comparisons using the BH method, with a statistical threshold of *q* < 0.05. Crucially, these analyses were performed only for the specific parent‒ offspring sex combinations and CM subtypes in which a significant direct effect on the LGI was initially identified.

Finally, in cases where significant associations were found between both (a) parental CM and parental psychosocial functioning and (b) parental psychosocial functioning and the offspring LGI, we performed a causal mediation analysis to test for indirect effects (74). This analysis was designed to determine whether parental psychosocial functioning mediated the relationship between parental CM subtypes and the offspring LGI. We implemented the models using the ’mediation’ package in R (75). This procedure estimates the average causal mediation effect (ACME; the indirect effect), the average direct effect (ADE), the total effect (TE), and the proportion mediated (PM). Statistical significance for each effect was determined by examining the 95% confidence intervals (CI), which were generated using a nonparametric bootstrapping procedure with 1,000 iterations using offspring age and CTQ total score as covariates. An effect was considered significant if its 95% CI did not include zero.

All the statistical analyses were performed in R version 4.3.1 (76).

## 3. RESULTS

### 3.1. Participant Characteristics

Following rigorous screening, two parent–offspring trios were excluded because of offspring neurological disorders and suboptimal brain imaging quality. Additionally, two fathers who exceeded the upper age limit for inclusion were excluded. Finally, 148 father–offspring dyads (60 sons and 88 daughters) and 150 mother–offspring dyads (60 sons and 90 daughters) were included in this study. The demographic characteristics of the sample are summarized in Table 1.

**Table 1.** Participant characteristics.

|  | Fathers with sons |  |  |  | Fathers with daughters |  |  |  | Mothers with sons |  |  |  | Mothers with daughters |  |  |  | Son |  |  |  | Daughter |  |  |  |
| --- | --- | --- | --- | --- | --- | --- | --- | --- | --- | --- | --- | --- | --- | --- | --- | --- | --- | --- | --- | --- | --- | --- | --- | --- |
|  | Mean | SD | Min | Max | Mean | SD | Min | Max | Mean | SD | Min | Max | Mean | SD | Min | Max | Mean | SD | Min | Max | Mean | SD | Min | Max |
| age | 52.28 | 5.89 | 41.00 | 65.00 | 54.72 | 6.08 | 43.00 | 65.00 | 50.98 | 5.07 | 41.00 | 63.00 | 53.51 | 5.52 | 41.00 | 64.00 | 19.27 | 3.47 | 15.00 | 33.00 | 22.17 | 6.71 | 15.00 | 38.00 |
| EA | 6.32 | 2.40 | 5.00 | 18.00 | 6.14 | 2.27 | 5.00 | 16.00 | 7.52 | 3.78 | 5.00 | 22.00 | 7.17 | 3.70 | 5.00 | 23.00 | 6.13 | 1.82 | 5.00 | 13.00 | 7.31 | 3.00 | 5.00 | 17.00 |
| PA | 5.48 | 1.19 | 5.00 | 11.00 | 5.41 | 1.47 | 5.00 | 16.00 | 5.55 | 1.76 | 5.00 | 15.00 | 5.41 | 1.36 | 5.00 | 15.00 | 5.07 | 0.31 | 5.00 | 7.00 | 5.18 | 0.59 | 5.00 | 9.00 |
| SA | 5.07 | 0.41 | 5.00 | 8.00 | 5.05 | 0.33 | 5.00 | 8.00 | 5.32 | 0.98 | 5.00 | 10.00 | 5.48 | 1.55 | 5.00 | 13.00 | 5.00 | 0.00 | 5.00 | 5.00 | 5.09 | 0.57 | 5.00 | 10.00 |
| EN | 12.83 | 3.88 | 5.00 | 24.00 | 12.63 | 4.64 | 5.00 | 25.00 | 12.72 | 5.13 | 5.00 | 25.00 | 11.68 | 5.17 | 5.00 | 24.00 | 10.62 | 4.07 | 5.00 | 23.00 | 10.18 | 4.54 | 5.00 | 25.00 |
| PN | 7.88 | 2.56 | 5.00 | 14.00 | 8.23 | 2.63 | 5.00 | 16.00 | 7.17 | 2.33 | 5.00 | 16.00 | 7.33 | 2.64 | 5.00 | 16.00 | 6.92 | 1.90 | 5.00 | 12.00 | 6.07 | 1.44 | 5.00 | 13.00 |
| CTQ total | 37.58 | 7.74 | 25.00 | 66.00 | 37.45 | 8.66 | 25.00 | 69.00 | 38.27 | 10.57 | 25.00 | 78.00 | 37.07 | 10.99 | 25.00 | 75.00 | 33.73 | 5.92 | 25.00 | 50.00 | 33.82 | 8.28 | 25.00 | 63.00 |
| K6 | 4.23 | 4.14 | 0.00 | 20.00 | 2.92 | 3.26 | 0.00 | 13.00 | 4.50 | 4.20 | 0.00 | 19.00 | 4.01 | 3.79 | 0.00 | 20.00 | - | - | - | - | - | - | - | - |
| AVO | 14.43 | 5.63 | 6.00 | 28.00 | 14.63 | 7.21 | 5.00 | 38.00 | 14.52 | 7.33 | 6.00 | 36.00 | 14.53 | 7.28 | 6.00 | 38.00 | - | - | - | - | - | - | - | - |
| ANX | 6.53 | 3.49 | 3.00 | 15.00 | 6.05 | 3.46 | 3.00 | 15.00 | 3.20 | 3.70 | 3.00 | 21.00 | 6.99 | 4.29 | 3.00 | 21.00 | - | - | - | - | - | - | - | - |

SD, standard deviation; EA, emotional abuse; PA, physical abuse; SA, sexual abuse; EN, emotional neglect; PN, physical neglect; CTQ, childhood trauma questionnaire; K6, Kessler 6; AVO, attachment-related avoidance; and ANX, attachment-related anxiety.

1. Participant characteristics onnaire; K6, Kessler 6; AVO, attachment-related avoidance; and ANX, attachment-related anxiety.

### 3.2. Associations between maternal/paternal CM and the offspring LGI

With respect to sons, maternal EA was significantly negatively associated with the LGI. This association was observed in several left-hemisphere regions, including the pre- and postcentral gyri and the paracentral region (Table 2, Figure 1). The other maternal CM subtypes, including PA, SA, EN, and PN, did not show any significant associations with the offspring LGI in any region. For daughters, no significant associations were found between any of the maternal CM subtypes and LGI.

**Figure 1.**
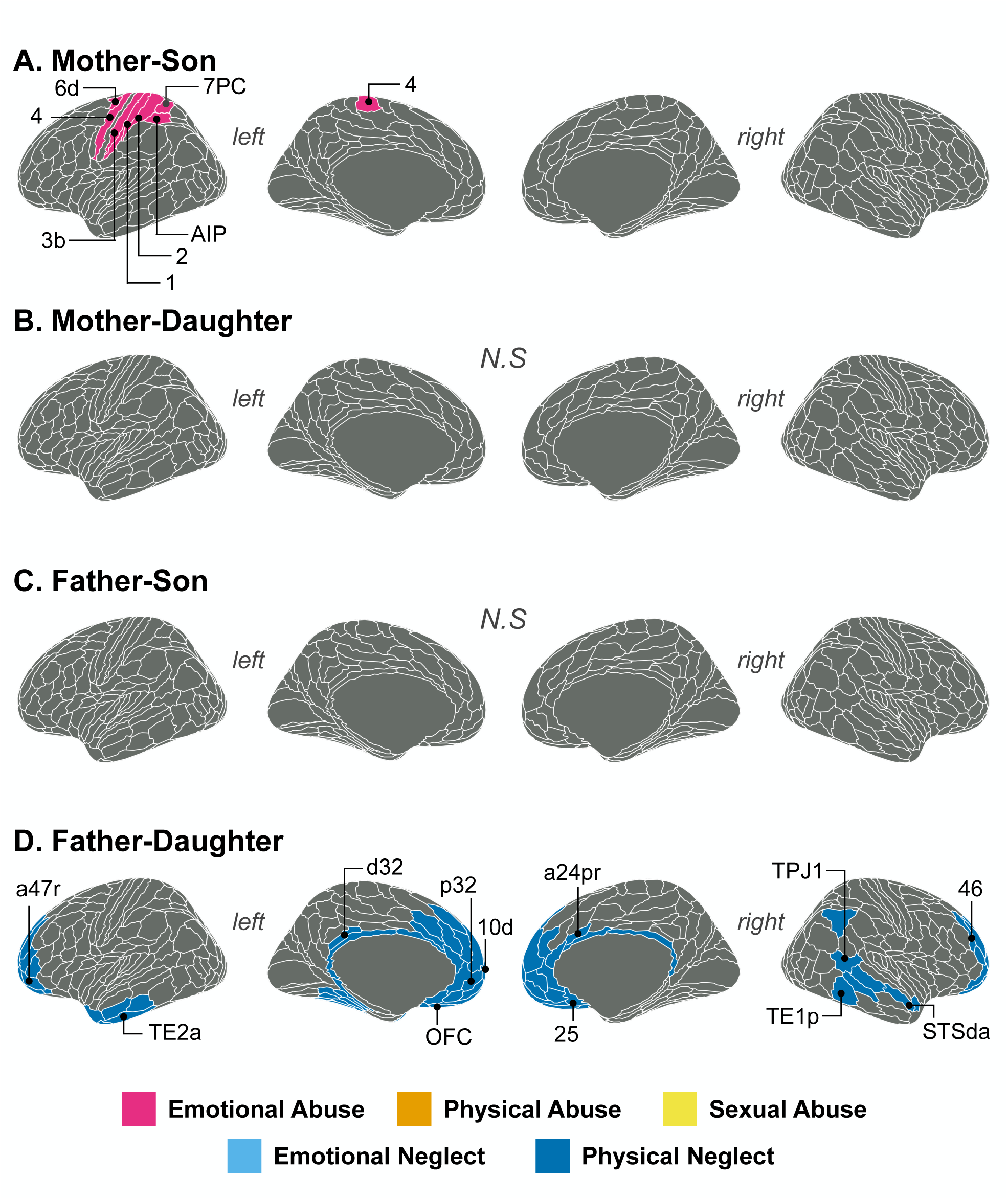
The associations between parental CM and offspring LGI Brain regions that the associations between parental CM and offspring LGI were statistically significant (the FDR-corrected *p*-values (*q*-values) < 0.05) are shown. Each brain region was colored according to maltreatment subtypes: pink, emotional abuse; orange, physical abuse; yellow, sexual abuse; light blue, emotional neglect; dark blue, physical neglect. The names of the brain regions followed the notation in Glasser et al (72). (A) The association between maternal CM and the LGI in sons, (B) the association between maternal CM and the LGI in daughters, (C) the association between paternal CM and the LGI in sons, (D) the association between paternal CM and the LGI in daughters. This figure was created using the ggseg plotting tool (77).

**Table 2.**
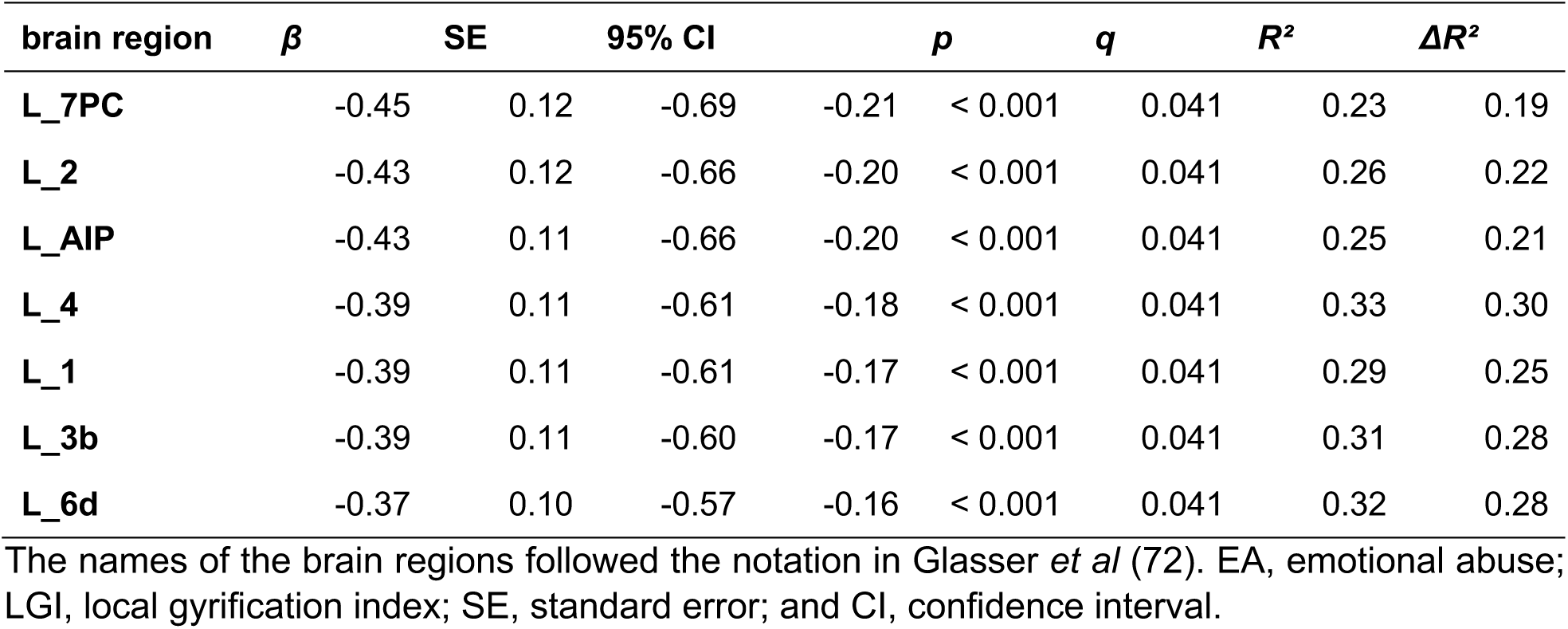
Result of maternal EA associated with the LGI in sons.

The analysis of paternal CM revealed a significant negative association between PN and the offspring LGI. This effect was specific to daughters and was observed in several regions, including the bilateral prefrontal cortex, cingulate cortex, and parts of the temporal cortex (Table 3, Figure 2). No other paternal CM subtypes were significantly associated with LGI in daughters, and no significant associations were found between any paternal CM subtype and the LGI in sons.

**Table 3.** Results of paternal PN associated with the LGI in daughters.

| brain region | $\beta$ | SE | 95% CI | | $p$ | $q$ | $R^2$ | $\Delta R^2$ |
| --- | --- | --- | --- | --- | --- | --- | --- | --- |
| L_10d | -0.40 | 0.10 | -0.60 | -0.21 | < 0.001 | 0.0094 | 0.21 | 0.18 |
| L_p32 | -0.40 | 0.10 | -0.59 | -0.21 | < 0.001 | 0.0094 | 0.21 | 0.18 |
| L_9m | -0.39 | 0.10 | -0.58 | -0.20 | < 0.001 | 0.0094 | 0.22 | 0.19 |
| L_a24 | -0.39 | 0.10 | -0.58 | -0.19 | < 0.001 | 0.0094 | 0.21 | 0.18 |
| L_10v | -0.39 | 0.10 | -0.59 | -0.18 | < 0.001 | 0.015 | 0.17 | 0.14 |
| L_10r | -0.38 | 0.10 | -0.58 | -0.18 | < 0.001 | 0.015 | 0.17 | 0.15 |
| L_10pp | -0.38 | 0.10 | -0.59 | -0.17 | < 0.001 | 0.015 | 0.16 | 0.13 |
| L_d32 | -0.38 | 0.093 | -0.56 | -0.19 | < 0.001 | 0.0094 | 0.21 | 0.19 |
| L_p24 | -0.37 | 0.094 | -0.56 | -0.19 | < 0.001 | 0.0094 | 0.20 | 0.17 |
| L_TE2a | -0.37 | 0.11 | -0.59 | -0.15 | 0.0012 | 0.015 | 0.12 | 0.092 |
| R_p32 | -0.37 | 0.10 | -0.57 | -0.16 | < 0.001 | 0.015 | 0.20 | 0.17 |
| R_10r | -0.37 | 0.10 | -0.57 | -0.16 | < 0.001 | 0.015 | 0.18 | 0.15 |
| R_10v | -0.36 | 0.10 | -0.57 | -0.15 | < 0.001 | 0.015 | 0.18 | 0.15 |
| R_a24 | -0.35 | 0.10 | -0.56 | -0.15 | < 0.001 | 0.015 | 0.18 | 0.15 |
| L_OFC | -0.35 | 0.10 | -0.56 | -0.14 | 0.0011 | 0.015 | 0.15 | 0.12 |
| R_9m | -0.35 | 0.10 | -0.54 | -0.15 | < 0.001 | 0.015 | 0.21 | 0.18 |
| L_9a | -0.35 | 0.10 | -0.54 | -0.15 | < 0.001 | 0.015 | 0.18 | 0.15 |
| L_TGv | -0.35 | 0.11 | -0.57 | -0.13 | 0.0024 | 0.021 | 0.13 | 0.10 |
| R_25 | -0.35 | 0.11 | -0.57 | -0.12 | 0.0027 | 0.023 | 0.13 | 0.10 |
| L_s32 | -0.35 | 0.10 | -0.55 | -0.14 | 0.0011 | 0.015 | 0.16 | 0.13 |
| R_9p | -0.35 | 0.10 | -0.53 | -0.16 | < 0.001 | 0.015 | 0.26 | 0.24 |
| L_a32pr | -0.34 | 0.094 | -0.53 | -0.16 | < 0.001 | 0.015 | 0.22 | 0.19 |
| L_25 | -0.34 | 0.10 | -0.55 | -0.14 | 0.0014 | 0.016 | 0.14 | 0.11 |
| L_a10p | -0.34 | 0.10 | -0.53 | -0.15 | < 0.001 | 0.015 | 0.17 | 0.14 |
| R_46 | -0.34 | 0.10 | -0.53 | -0.14 | < 0.001 | 0.015 | 0.22 | 0.19 |
| L_11l | -0.33 | 0.10 | -0.53 | -0.13 | 0.0013 | 0.015 | 0.15 | 0.12 |
| R_10pp | -0.33 | 0.10 | -0.53 | -0.13 | 0.0016 | 0.017 | 0.17 | 0.14 |
| R_s32 | -0.33 | 0.11 | -0.54 | -0.12 | 0.0027 | 0.023 | 0.16 | 0.13 |
| R_10d | -0.33 | 0.10 | -0.53 | -0.13 | 0.0013 | 0.015 | 0.19 | 0.16 |
| R_9a | -0.33 | 0.10 | -0.52 | -0.14 | < 0.001 | 0.015 | 0.23 | 0.20 |
| R_p24 | -0.33 | 0.10 | -0.52 | -0.14 | 0.0011 | 0.015 | 0.19 | 0.16 |
| R_9-46d | -0.33 | 0.093 | -0.51 | -0.14 | < 0.001 | 0.015 | 0.24 | 0.22 |
| L_TF | -0.33 | 0.10 | -0.53 | -0.12 | 0.0021 | 0.020 | 0.12 | 0.084 |
| L_p10p | -0.33 | 0.10 | -0.52 | -0.14 | < 0.001 | 0.015 | 0.18 | 0.15 |
| L_a24pr | -0.33 | 0.094 | -0.51 | -0.14 | < 0.001 | 0.015 | 0.22 | 0.20 |
| R_pOFC | -0.32 | 0.11 | -0.55 | -0.10 | 0.0049 | 0.033 | 0.11 | 0.083 |
| R_OFC | -0.32 | 0.11 | -0.53 | -0.11 | 0.0039 | 0.029 | 0.15 | 0.12 |
| L_pOFC | -0.32 | 0.11 | -0.53 | -0.11 | 0.0034 | 0.026 | 0.11 | 0.078 |
| L_p47r | -0.32 | 0.11 | -0.53 | -0.10 | 0.0045 | 0.031 | 0.13 | 0.10 |
| R_a10p | -0.31 | 0.10 | -0.51 | -0.12 | 0.0018 | 0.018 | 0.20 | 0.17 |
| R_33pr | -0.31 | 0.095 | -0.50 | -0.12 | 0.0016 | 0.017 | 0.19 | 0.16 |
| L_a47r | -0.31 | 0.10 | -0.51 | -0.10 | 0.0038 | 0.028 | 0.13 | 0.10 |
| R_d32 | -0.31 | 0.10 | -0.50 | -0.11 | 0.0023 | 0.021 | 0.20 | 0.17 |
| L_RSC | -0.30 | 0.10 | -0.50 | -0.10 | 0.0032 | 0.025 | 0.18 | 0.15 |
| L_TE1a | -0.30 | 0.11 | -0.51 | -0.090 | 0.0059 | 0.037 | 0.10 | 0.072 |
| L_TGd | -0.30 | 0.11 | -0.51 | -0.085 | 0.0067 | 0.039 | 0.12 | 0.094 |
| R_STSvp | -0.30 | 0.094 | -0.49 | -0.11 | 0.0022 | 0.020 | 0.14 | 0.10 |
| L_VMV3 | -0.30 | 0.087 | -0.47 | -0.12 | < 0.001 | 0.015 | 0.18 | 0.15 |
| R_p10p | -0.30 | 0.092 | -0.48 | -0.11 | 0.0018 | 0.018 | 0.21 | 0.18 |
| R_a9-46v | -0.29 | 0.10 | -0.49 | -0.10 | 0.0032 | 0.025 | 0.16 | 0.13 |
| L_VMV2 | -0.29 | 0.087 | -0.46 | -0.12 | 0.0013 | 0.015 | 0.18 | 0.15 |
| R_RSC | -0.29 | 0.10 | -0.48 | -0.093 | 0.0044 | 0.031 | 0.12 | 0.084 |
| L_VMV1 | -0.29 | 0.088 | -0.46 | -0.11 | 0.0016 | 0.017 | 0.18 | 0.15 |
| L_33pr | -0.28 | 0.10 | -0.48 | -0.092 | 0.0042 | 0.030 | 0.20 | 0.17 |
| L_8BM | -0.28 | 0.093 | -0.47 | -0.10 | 0.0032 | 0.025 | 0.21 | 0.18 |
| R_TPOJ1 | -0.28 | 0.10 | -0.48 | -0.086 | 0.0054 | 0.035 | 0.14 | 0.11 |
| L_a9-46v | -0.28 | 0.10 | -0.47 | -0.086 | 0.0050 | 0.033 | 0.16 | 0.13 |
| R_TE1p | -0.28 | 0.10 | -0.48 | -0.067 | 0.010 | 0.049 | 0.078 | 0.045 |
| L_TE1m | -0.27 | 0.10 | -0.48 | -0.067 | 0.010 | 0.049 | 0.088 | 0.055 |
| L_d23ab | -0.27 | 0.10 | -0.48 | -0.069 | 0.0094 | 0.047 | 0.16 | 0.13 |
| R_STSda | -0.27 | 0.10 | -0.47 | -0.075 | 0.0074 | 0.042 | 0.12 | 0.084 |
| R_11l | -0.27 | 0.10 | -0.47 | -0.068 | 0.0094 | 0.047 | 0.16 | 0.13 |
| L_VVC | -0.27 | 0.092 | -0.45 | -0.086 | 0.0045 | 0.031 | 0.13 | 0.10 |
| L_47m | -0.27 | 0.10 | -0.46 | -0.074 | 0.0071 | 0.041 | 0.12 | 0.089 |
| L_PreS | -0.26 | 0.094 | -0.45 | -0.078 | 0.0060 | 0.037 | 0.12 | 0.089 |
| R_STSdp | -0.26 | 0.10 | -0.46 | -0.071 | 0.0080 | 0.044 | 0.11 | 0.083 |
| **L_p32pr** | -0.26 | 0.094 | -0.45 | -0.076 | 0.0065 | 0.038 | 0.22 | 0.19 |
| **R_a47r** | -0.26 | 0.10 | -0.46 | -0.068 | 0.0086 | 0.044 | 0.15 | 0.12 |
| **L_9-46d** | -0.26 | 0.10 | -0.46 | -0.069 | 0.0085 | 0.044 | 0.17 | 0.14 |
| **R_STGa** | -0.26 | 0.094 | -0.45 | -0.075 | 0.0065 | 0.038 | 0.11 | 0.075 |
| **L_PHA1** | -0.26 | 0.092 | -0.44 | -0.077 | 0.0059 | 0.037 | 0.13 | 0.10 |
| **R_a24pr** | -0.25 | 0.10 | -0.44 | -0.061 | 0.0104 | 0.049 | 0.17 | 0.14 |
| **R_8BL** | -0.25 | 0.093 | -0.44 | -0.067 | 0.0084 | 0.044 | 0.22 | 0.19 |
| **L_PHA3** | -0.25 | 0.092 | -0.43 | -0.067 | 0.0078 | 0.043 | 0.12 | 0.088 |
| **R_PFm** | -0.25 | 0.092 | -0.43 | -0.065 | 0.0083 | 0.044 | 0.17 | 0.14 |
| **L_PHA2** | -0.24 | 0.092 | -0.42 | -0.059 | 0.010 | 0.049 | 0.11 | 0.083 |
| **L_9p** | -0.24 | 0.090 | -0.41 | -0.057 | 0.010 | 0.049 | 0.18 | 0.15 |

The names of the brain regions followed the notation in Glasser *et al* (72). PN, physical neglect; LGI, local gyrification index; SE, standard error; and CI, confidence interval.

### 3.3. Associations between maternal/paternal CM and own psychosocial functioning

We subsequently examined the association between maternal EA—the subtype that was significantly linked to the LGI among sons—and the mother’s own psychosocial functioning. No significant associations were found with any of the psychosocial metrics (K6, *p* = 0.056, *q* = 0.083; AVO, *p* = 0.083, *q* = 0.083; ANX, *p* = 0.048, *q* = 0.083).

Similarly, we examined the association between paternal PN—the subtype linked to the LGI among daughters—and the father’s own psychosocial functioning. No significant associations were detected (K6, *p* = 0.091, *q* = 0.22; AVO, *p* = 0.15, *q* = 0.22; ANX, *p* = 0.77, *q* = 0.77).

### 3.4. Associations between maternal/paternal psychosocial functioning and the offspring LGI

The analysis of the relationship between maternal psychosocial functioning and the LGI among sons yielded one significant finding. Among the three maternal variables, only ANX was significantly associated with the LGI in all seven of the regions previously identified in the maternal EA analysis for sons (Table 4). In contrast, neither AVO nor K6 was significantly associated.

**Table 4.**
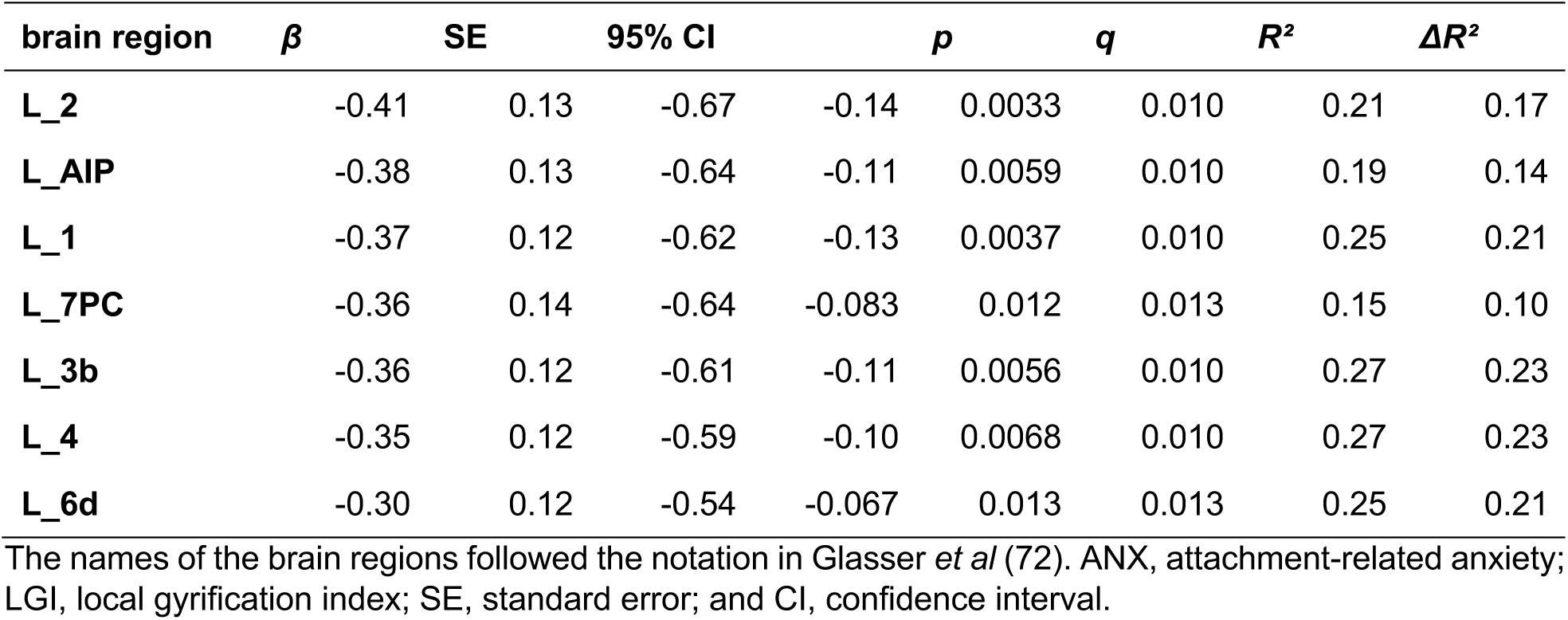
Results of maternal ANX associated with the LGI in sons.

None of the paternal psychosocial functioning metrics were significantly associated with the LGI in any of the brain regions in daughters.

### 3.5. Mediation analysis

Although the direct association between maternal EA and ANX did not reach statistical significance, we conducted a causal mediation analysis as a supplementary investigation. The analysis did not yield a significant mediation effect, indicating that the relationship between maternal EA and the LGI in sons is likely independent of the mother’s ANX.

## 4. DISCUSSION

In the present study, we demonstrated an intergenerational association between parental CM and the LGI among adolescent and young adult offspring. More specifically, our findings revealed a distinct sex-specific pattern: maternal EA was negatively associated with the LGI in sons, whereas paternal PN was negatively associated with the LGI in daughters. We also found that these associations were not mediated by parental psychosocial functioning, suggesting the influence of an early biological transmission pathway.

### 4.1. Maternal childhood maltreatment and offspring cortical gyrification

A study by Mareckova *et al*. (2020) demonstrated that maternal prenatal stress is specifically and negatively correlated with the LGI in sons (51). Our findings show some regional overlap with those of Mareckova *et al*. (2020). Specifically, the associations they reported in the left superior frontal gyrus (extending anterior to the central sulcus) and the medial paracentral region are partially consistent with our results. In animal models, maternal prenatal stress has been shown to induce numerous changes in the cerebral cortex of offspring, including alterations in dendritic morphology and synaptic connectivity (78), a reduction in neural progenitor cells and neurogenesis (79), and a disruption of the excitation/inhibition balance resulting from the dysregulated expression of glutamatergic genes (80). Notably, the pericentral sulcus region, where we found a significant association, is an area that develops early in the fetal period and shows low variability thereafter (81). Furthermore, it has been reported in human studies that this same region is affected by very preterm birth and is also associated with the intelligence quotient (IQ) and psychiatric symptoms at age 30 (52). It is therefore conceivable that the observed alterations in gyrification resulted from a complex interplay of various maternal factors surrounding the perinatal period, which may represent the underlying molecular mechanism of our findings.

In particular, the study by Mareckova *et al*. (2020) investigated prenatal stress, not CM. However, previous studies have indicated that these two factors are closely associated. Souch *et al*. (2022) reported that maternal CM was associated with adverse perinatal outcomes, including perinatal depression (7). Additionally, these associations were found to be particularly consistent for the EA and SA subscales. Threat exposure, such as EA and SA, has been robustly associated with the fronto-amygdala circuit, which plays a key role in emotional processes (82). These regions notably include the amygdala and the ventromedial prefrontal cortex, where both structural and functional alterations have been reported in individuals exposed to CM (3,4,83). According to a systematic review, alterations in these brain regions are consistently observed in offspring with a history of threat exposure. In contrast, such consistent alterations were not apparent in offspring who experienced deprivation (84). The fronto-amygdala circuit has been strongly implicated in human emotion regulation (85,86), and structural and functional abnormalities in these regions have been linked to various psychiatric disorders, including substance use disorder (87), schizophrenia (88), and anxiety disorders (89). Therefore, our findings suggest that the experiences of mothers with EA influence their emotion regulation processes, which in turn are associated with the development of the brains of their offspring.

Notably, our results indicated a sex-specific effect: the association was significant exclusively for sons and was not observed for daughters. These findings may be explained by the hypothesis that the effects of prenatal stress on offspring are sex specific. As mentioned previously, Mareckova *et al*. (2020) reported sex-specific effects of maternal prenatal stress on the offspring LGI (51). Although research specifically examining the sex-specific effects of maternal CM on offspring remains scarce (17), findings from the field of gestational biology indicate that maternal–fetal physiological processes differ depending on the sex of the fetus. The placenta, which is responsible for signaling from mother to fetus, responds to changes in maternal stress differently depending on the sex of the fetus (90). For example, research in animal models has shown that in response to prenatal stress, the inflammatory response is more robust in the male placenta than in the female placenta (91). Furthermore, the placental expression of O-linked-*N*- acetylglucosamine (O-GlcNAc) transferase (OGT), a gene on the X chromosome widely involved in regulating protein function, has been shown to be specifically lower in males and to be further reduced by prenatal stress (92,93). Furthermore, it has been suggested that male fetuses adopt a strategy of maintaining normal growth even in an adverse intrauterine environment, which may consequently increase their vulnerability to the severe effects of subsequent stressful events (94). Mothers have also been shown to be differentially affected by their pregnancy depending on the sex of their fetus. For instance, higher cortisol levels have been observed in mothers pregnant with male fetuses than in those pregnant with female fetuses (95). Furthermore, compared with mothers who carry female fetuses, mothers who carry male fetuses have a greater risk of developing postpartum depression (96–98). A potential biological mechanism for this association is the finding that estrogen levels, which are implicated in postpartum depression, are lower in mothers of male fetuses (99,100). In a study analyzing the sex-specific intergenerational effects of maternal ACEs, an indirect pathway in which perinatal maternal mental health mediated the effects on internalizing and externalizing problems among male offspring was found (101). Although this remains speculative, it is plausible that maternal history of CM led to an accumulation of insults during the prenatal period or earlier, which, through various physiological pathways, ultimately contributed to the observed LGI development specifically in their male offspring.

### 4.2. Paternal childhood maltreatment and offspring cortical gyrification

Addressing a significant gap in the literature, Karlsson and colleagues pioneered research on the effects of paternal CM on offspring brain development. Using data from the FinnBrain Birth Cohort, they investigated the effects of both paternal and maternal CM on the brain structure of neonates (32–34). The results of the present study differ from those of previous studies in terms of specific metrics and the developmental stage of the offspring, making direct comparisons of the results difficult. Nevertheless, our findings provide a novel contribution to the understanding of how paternal CM may be transmitted across generations to affect brain development.

In the present study, we demonstrated an association between paternal PN and the LGI in daughters during adolescence and young adulthood. This transmission mechanism might be explained by epigenetic alterations in the father’s sperm. Several previous studies in healthy human males have reported the presence of epigenetic alterations in the sperm of men who experienced early-life stress (10–12). Roberts *et al*. (2018) reported that childhood abuse was associated with DNA methylation (DNAm) in 12 distinct regions within the sperm of adult males. These genetic regions are broadly involved in pathways governing neural function, fat cell regulation, and immune response (10). In addition, another research group reported reduced levels of several miRNAs from the miR-449/34 family in the sperm of adult men who had experienced multiple ACEs. Moreover, a similar trend was observed in the sperm of male mice exposed to chronic social instability stress. This effect was shown to persist into the early embryos sired by these mice and was even present in the sperm of their male offspring (11). Furthermore, these miRNA changes have been partially replicated in human studies by several different research groups (12,102). This convergence of evidence lends credence to the possibility that early-life stress induces epigenetic alterations in male sperm, which are then transmitted to the next generation. This hypothesis is further substantiated by the only prior study that has reported on the effects of paternal ACEs on offspring DNAm (103). In a study of 3- month-old infants, Merrill *et al*. (2021) performed an epigenome-wide analysis on the association between paternal ACEs and blood DNAm. They reported significant associations at CpG sites in eight regions related to reproductive and metabolic functions, suggesting the possibility that biomarkers linked to paternal trauma are already established early in the postnatal period (103). The association between paternal CM and the offspring LGI observed in the present study may therefore be underpinned by a mechanism of intergenerational transmission involving epigenetic changes in sperm.

The finding that only PN was associated with the offspring LGI in our analysis of paternal CM might be explained by sex differences in exposure and vulnerability to various types of CM. A meta-analysis revealed that females report SA at twice the rate of males (104). Furthermore, GMV in the hippocampus and postcentral gyrus has been reported to be associated with childhood neglect and deprivation, specifically among male victims (105,106). These findings suggest that males may be more sensitive than females to the effects of neglect (107). Additionally, in a meta-analysis of the effects of ACEs, PN was associated with both depression and anxiety in males, whereas no significant associations were found for these outcomes in females (108). Since prior research on CM subtypes such as neglect is limited (109), our interpretation remains speculative. Nevertheless, we propose that the association found only with paternal PN may be related to a greater vulnerability of males to this type of adversity.

A key finding of our study was that the impact of paternal CM was evident exclusively in daughters. The interpretation of the underlying mechanism is challenging, given the scarcity of human studies that have analyzed paternal influences separately for male and female offspring. Nevertheless, the results from animal models may offer some potential explanations. Consistent with this, several studies in mice have suggested that the effects of paternal environmental exposure can be transmitted in a sex-specific manner, selectively affecting female offspring (110–114). Franklin *et al*. (2010) reported that female offspring of male mice exposed to unpredictable maternal separation exhibited depressive-like behaviors and altered DNAm in brain genes related to stress resistance, emotion, and depression (110). Additionally, He *et al*. (2016) reported that the female offspring of males exposed to social instability stress during adolescence exhibited increased anxiety, elevated cortisol levels, and decreased hippocampal mRNA expression of both glucocorticoid receptors and brain-derived neurotrophic factor (BDNF) (112). Potential mechanisms underlying these sex-specific effects include differences in sex chromosomes and sex hormones, differential epigenetic changes in the placenta, and sex-dependent variations in nutrient transport via the placenta (115–117). While such mechanisms might explain our findings, there are considerable cross-species differences in gestational biology, with fundamental distinctions between humans and rodent models (118). Therefore, further research in humans is warranted.

In the present study, the association between paternal CM and the offspring LGI was observed across a wide range of brain regions, extending from the medial and lateral prefrontal cortex to the cingulate gyrus and parts of the temporal lobe. Karlsson *et al*. (2020) revealed that paternal CM affects the offspring’s corpus callosum, a structure that reflects the capacity for information transfer between the left and right cerebral hemispheres (32). The corpus callosum is known to be particularly vulnerable to the effects of CM, with robust reports suggesting that morphological changes in this structure may be linked to variations in problem-solving skills and IQ (83). The corpus callosum and cingulate gyrus are not only anatomically adjacent, with the corpus callosum located directly inferior to the gyrus, but they are also functionally connected via the cingulum bundle (119,120). Thus, some of our results corroborate previous findings on the impact of CM on the corpus callosum. Moreover, the anterior cingulate cortex and prefrontal cortex are known to function as crucial hubs for emotion regulation and are implicated in various psychiatric disorders (121). Consistent with this, numerous studies have reported associations between the structure of these regions and clinical symptoms, particularly during the developmental period spanning adolescence and young adulthood (122–125). Furthermore, a growing body of literature suggests that a father’s own history of adversity increases the risk of psychiatric disorders and problem behaviors among his offspring (41,103,126–131). Although the present study did not directly examine the link between the offspring’s psychiatric symptoms and their brain structure, the LGI alterations associated with paternal CM may represent a potential neural substrate for the intergenerational transmission of psychiatric risk.

### 4.3. Maternal and paternal childhood maltreatment and own behaviors

To further elucidate the mechanisms through which paternal and maternal CM are transmitted to affect the offspring LGI, we analyzed its associations with parental psychosocial functions. In the maternal analysis, while we found significant associations between both EA and ANX with the LGI of sons, the indirect effect was not statistically significant. Furthermore, in the paternal analysis, neither the association between PN and paternal psychosocial functioning nor the association between their psychosocial functioning and the offspring LGI was significant. These findings suggest that the influence of parental CM on the offspring LGI is likely independent of parental psychosocial functions.

Hypothesizing that the intergenerational transmission of the impact of parental CM is established prenatally, the present study analyzed the LGI as a potential biomarker. Although establishing a causal link between an exposure and a phenotype is extremely challenging in human studies, the use of neuroimaging can help mitigate this limitation (32). As previously mentioned, the LGI is a neurodevelopmental metric that begins its formation prenatally and is considered a relatively stable metric throughout development. Thus, in studying the intergenerational transmission between parental CM and offspring brain development, the LGI offers a distinct advantage. Because it is a relatively stable marker established early in life, it substantially minimizes the confounding influence of postnatal environmental factors (e.g., parenting and lifestyle factors) compared with more plastic metrics such as GMV or cortical thickness. Consistent with this hypothesis, although our analyses revealed some significant associations with parental marital relationships and mental health, we ultimately did not detect any factors that mediated the relationship between parental CM and the offspring LGI. Specifically, a mother’s own history of CM has been shown to increase her risk for various adverse outcomes, including her own prenatal depression, perinatal complications, and marital discord (7). Notably, we did not collect data on these maternal factors, such as perinatal outcomes and marital discord, from the offspring’s early childhood. Therefore, while it is plausible that these factors contributed to our findings, we were unable to analyze this potential pathway in the present study. Despite the aforementioned limitations, the primary finding of the present study was an association between maternal and paternal CM and the offspring LGI that was not mediated by the current mental health or marital relationships of the parents. This finding supports the hypothesis that the intergenerational transmission of parental CM effects is established prenatally or even earlier.

### 4.4. Limitations

The present study has several limitations. First, the sample of offspring was characterized by an imbalanced sex ratio, with a larger number of females than males. Consequently, our analyses of sex-specific effects may have been underpowered, potentially limiting our ability to detect true effects in the smaller male subgroup. Second, parental CM was retrospectively assessed using a self-report measure, which is susceptible to recall bias and potential overreporting. However, both prospective and retrospective assessments of CM have demonstrated associations with developmental outcomes in the same direction, although the effect sizes may differ slightly (37,132). Third, the study’s cross-sectional design prevents the clarification of several key aspects. Specifically, we could not determine the prospective associations with occurrences from the fetal or early childhood periods, nor could we trace the developmental trajectory of the LGI effects from the fetal period to the present.

## 5. CONCLUSION

In conclusion, our study of biological parent–offspring trios provides evidence that CM experienced by parents is associated with offspring brain structure, as measured by the LGI. Furthermore, we discovered distinct sex-specific transmission pathways: paternal PN was associated with the LGI in daughters, whereas maternal EA was associated with the LGI in sons. These findings collectively strengthen the hypothesis that the neurodevelopmental consequences of parental CM are transmitted intergenerationally through early biological mechanisms, potentially established prenatally or even prior to conception.

## ACKNOWLEDGMENTS

This research received funding from a Grant-in-Aid for the Japan Society for the Promotion of Science (JSPS) Fellows (Grant No. 23KJ0220) and Japan Science and Technology agency Support for Pioneering Research Initiated by the Next Generation (JST SPRING) (Grant No. JPMJSP2114) to RY. IM was supported by a Grant-in-Aid for Young Scientists (Grant No. 22K13809), Grant-in-Aid for Transformative Research Areas (A) (Grant Nos. 22H05209 and 24H00896), and Grant-in-Aid for Scientific Research (B) (Grant No. 23K27258), provided by the Ministry of Education, Culture, Sports, and Technology (MEXT) and JSPS. Additionally, it was supported by the Program for the Creation of Interdisciplinary Research from the Frontier Research Institute for Interdisciplinary Sciences, Tohoku University, and a research grant from the Intelligent Cosmos Academic Foundation awarded to IM. Further support was provided by a Grant-in-Aid for Scientific Research (B) (Grant No. 19H04211) from MEXT to YT. The preprocessing and statistical analysis of the neuroimaging data were funded by JSPS KAKENHI Grant Number JP22H04926 and the Grant-in-Aid for Transformative Research Areas— Platforms for Advanced Technologies and Research Resources “Advanced Bioimaging Support.” We wish to express our deepest thanks to the participants for their willingness to participate in this study. Special thanks are directed to Yukiko Suginome, Saeko Hoshi, Chieko Miura, Maiko Chiba, Junko Kato, Misaki Abe, and Shuzo Yamamoto for their invaluable support. We are also grateful to Dr. Ryosuke Kimura, Dr. Tadashi Imanishi, Ayaka Uchiyama, Kanna Oyama, Mihiro Koizumi, Takumi Uchiyama, Yuka Aoki, Fumiaki Nitta, Yuka Hatayama, Megumi Kato, Maiko Suenaga, Jun Nomura, Ryuhei Ohgi, Ruri Takahashi, Yuto Tanaka, Saki Uchida, Ruriko Igarashi, and all other staff for their contributions to data collection. Additionally, we extend our gratitude to Drs. Kiyotaka Nemoto and Takuya Hayashi for their support with the neuroimaging analysis.

## AUTHOR CONTRIBUTIONS

R.Y. and I.M. designed the research. R.Y. and I.M. collected the data. R.Y. and I.M. analyzed the data. R.Y., I.M., and Y.T. acquired funding. Y.T. supervised this study. R.Y. and I.M. wrote and edited the article. All the authors read and approved the final version of the article.

## DECLARATION OF GENERATIVE AI AND AI-ASSISTED TECHNOLOGIES IN THE WRITING PROCESS

During the preparation of this work, we used ChatGPT to generate R scripts for statistical analysis. After using this tool, we reviewed and edited the content as needed, and we take full responsibility for the content of the publication.

## REFERENCES

1. Baldwin JR, Wang B, Karwatowska L, Schoeler T, Tsaligopoulou A, Munafò MR, Pingault J-B (2023): Childhood maltreatment and mental health problems: A systematic review and meta-analysis of quasi-experimental studies. Am J Psychiatry 180: 117–126.

2. Daníelsdóttir HB, Aspelund T, Shen Q, Halldorsdottir T, Jakobsdóttir J, Song H, et al. (2024): Adverse childhood experiences and adult mental health outcomes. JAMA Psychiatry 81: 586–594.

3. Nelson CA, Sullivan EF, Valdes V (2025): Early adversity alters brain architecture and increases susceptibility to mental health disorders. Nat Rev Neurosci 1–15.

4. Vannucci A, Fields A, Hansen E, Katz A, Kerwin J, Tachida A, et al. (2023): Interpersonal early adversity demonstrates dissimilarity from early socioeconomic disadvantage in the course of human brain development: A meta-analysis. Neurosci Biobehav Rev 150: 105210.

5. Hosseini-Kamkar N, Varvani Farahani M, Nikolic M, Stewart K, Goldsmith S, Soltaninejad M, et al. (2023): Adverse life experiences and brain function: A meta-analysis of functional magnetic resonance imaging findings: A meta-analysis of functional magnetic resonance imaging findings. JAMA Netw Open 6: e2340018.

6. McKay MT, Kilmartin L, Meagher A, Cannon M, Healy C, Clarke MC (2022): A revised and extended systematic review and meta-analysis of the relationship between childhood adversity and adult psychiatric disorder. J Psychiatr Res 156: 268–283.

7. Souch AJ, Jones IR, Shelton KHM, Waters CS (2022): Maternal childhood maltreatment and perinatal outcomes: A systematic review. J Affect Disord 302: 139–159.

8. Lange BCL, Callinan LS, Smith MV (2019): Adverse childhood experiences and their relation to parenting stress and parenting practices. Community Ment Health J 55: 651–662.

9. Condon EM, Dettmer A, Baker E, McFaul C, Stover CS (2022): Early life adversity and males: Biology, behavior, and implications for fathers’ parenting. Neurosci Biobehav Rev 135: 104531.

10. Roberts AL, Gladish N, Gatev E, Jones MJ, Chen Y, MacIsaac JL, et al. (2018): Exposure to childhood abuse is associated with human sperm DNA methylation. Transl Psychiatry 8: 194.

11. Dickson DA, Paulus JK, Mensah V, Lem J, Saavedra-Rodriguez L, Gentry A, et al. (2018): Reduced levels of miRNAs 449 and 34 in sperm of mice and men exposed to early life stress. Transl Psychiatry 8: 101.

12. Tuulari JJ, Bourgery M, Iversen J, Koefoed TG, Ahonen A, Ahmedani A, et al. (2025): Exposure to childhood maltreatment is associated with specific epigenetic patterns in sperm. Mol Psychiatry 1– 10.

13. Skjothaug T, Smith L, Wentzel-Larsen T, Moe V (2015): Prospective fathers’ adverse childhood experiences, pregnancy-related anxiety, and depression during pregnancy: Prospective fathers’ adverse childhood experiences. Infant Ment Health J 36: 104–113.

14. Skjothaug T, Smith L, Wentzel-Larsen T, Moe V (2018): Does fathers’ prenatal mental health bear a relationship to parenting stress at 6 months? Infant Ment Health J 39: 537–551.

15. Vaillancourt-Morel M-P, Bussières È-L, Nolin M-C, Daspe M-È (2024): Partner effects of childhood maltreatment: A systematic review and meta-analysis. Trauma Violence Abuse 25: 1150–1167.

16. Narayan AJ, Lieberman AF, Masten AS (2021): Intergenerational transmission and prevention of adverse childhood experiences (ACEs). Clin Psychol Rev 85: 101997.

17. Moog NK, Heim CM, Entringer S, Simhan HN, Wadhwa PD, Buss C (2022): Transmission of the adverse consequences of childhood maltreatment across generations: Focus on gestational biology. Pharmacol Biochem Behav 215: 173372.

18. Racine N, Deneault A-A, Thiemann R, Turgeon J, Zhu J, Cooke J, Madigan S (2023): Intergenerational transmission of parent adverse childhood experiences to child outcomes: A systematic review and meta-analysis. Child Abuse Negl 106479.

19. Racine N, Bellis MA, Madigan S (2024): An introduction to twenty-five years of adverse childhood experiences: A special issue. Child Abuse Negl 107224.

20. Flint J, Timpson N, Munafò M (2014): Assessing the utility of intermediate phenotypes for genetic mapping of psychiatric disease. Trends Neurosci 37: 733–741.

21. Liu C, Gershon ES (2024): Endophenotype 2.0: updated definitions and criteria for endophenotypes of psychiatric disorders, incorporating new technologies and findings. Transl Psychiatry 14: 502.

22. Moog NK, Entringer S, Rasmussen JM, Styner M, Gilmore JH, Kathmann N, et al. (2018): Intergenerational Effect of Maternal Exposure to Childhood Maltreatment on Newborn Brain Anatomy. Biol Psychiatry 83: 120–127.

23. Hendrix CL, Dilks DD, McKenna BG, Dunlop AL, Corwin EJ, Brennan PA (2021): Maternal Childhood Adversity Associates With Frontoamygdala Connectivity in Neonates. Biol Psychiatry Cogn Neurosci Neuroimaging 6: 470–478.

24. Zhang H, Wong T-Y, Broekman BFP, Chong Y-S, Shek LP, Gluckman PD, et al. (2021): Maternal Adverse Childhood Experience and Depression in Relation with Brain Network Development and Behaviors in Children: A Longitudinal Study. Cereb Cortex 31: 4233–4244.

25. Khoury JE, Ahtam B, Sisitsky M, Ou Y, Gagoski B, Enlow MB, et al. (2022): Maternal Childhood Maltreatment Is Associated With Lower Infant Gray Matter Volume and Amygdala Volume During the First Two Years of Life. Biological Psychiatry Global Open Science 2: 440–449.

26. Demers CH, Hankin BL, Hennessey E-MP, Haase MH, Bagonis MM, Kim SH, et al. (2022): Maternal adverse childhood experiences and infant subcortical brain volume. Neurobiology of Stress 21: 100487.

27. Lugo-Candelas C, Chang L, Dworkin JD, Aw N, Fields A, Reed H, et al. (2023): Maternal childhood maltreatment: associations to offspring brain volume and white matter connectivity. J Dev Orig Health Dis 14: 591–601.

28. van den Heuvel MI, Monk C, Hendrix CL, Hect J, Lee S, Feng T, Thomason ME (2023): Intergenerational Transmission of Maternal Childhood Maltreatment Prior to Birth: Effects on Human Fetal Amygdala Functional Connectivity. J Am Acad Child Adolesc Psychiatry. 10.1016/j.jaac.2023.03.020

29. Demers CH, Hankin BL, Haase MH, Todd E, Hoffman MC, Epperson CN, et al. (2024): Maternal adverse childhood experiences and infant visual-limbic white matter development. J Affect Disord 367: 49– 57.

30. Uy JP, Parks KC, Tan AP, Fortier MV, Meaney M, Chong YS, et al. (2025): Maternal childhood maltreatment, development of amygdala volume, and anxiety symptoms in offspring. J Am Acad Child Adolesc Psychiatry. 10.1016/j.jaac.2025.03.027

31. Lyons-Ruth K, Li FH, Khoury JE, Ahtam B, Sisitsky M, Ou Y, et al. (2023): Maternal childhood abuse versus neglect associated with differential patterns of infant brain development. Res Child Adolesc Psychopathol 51: 1919–1932.

32. Karlsson H, Merisaari H, Karlsson L, Scheinin NM, Parkkola R, Saunavaara J, et al. (2020): Association of Cumulative Paternal Early Life Stress With White Matter Maturation in Newborns. JAMA Netw Open 3: e2024832.

33. Tuulari JJ, Kataja E-L, Karlsson L, Karlsson H (2022, October 20): Associations of cumulative paternal and maternal childhood maltreament exposure with neonate brain anatomy. BioRxiv. p 2022.10.16.512276.

34. Tuulari JJ, Pulli EP, Kataja E-L, Perasto L, Lewis JD, Karlsson L, Karlsson H (2023, February 23): Parental childhood maltreatment associates with offspring left amygdala volume at early infancy. BioRxiv. 10.1101/2023.02.23.529799

35. Soubry A (2018): POHaD: why we should study future fathers. Environ Epigenet 4: dvy007.

36. Soubry A (2018): Epigenetics as a driver of developmental origins of health and disease: Did we forget the fathers? Bioessays 40: 1700113.

37. Zhang L, Mersky JP, Gruber AMH, Kim J-Y (2022): Intergenerational Transmission of Parental Adverse Childhood Experiences and Children’s Outcomes: A Scoping Review. Trauma Violence Abuse 15248380221126186.

38. Kaczkurkin AN, Raznahan A, Satterthwaite TD (2019): Sex differences in the developing brain: insights from multimodal neuroimaging. Neuropsychopharmacology 44: 71–85.

39. Lawrence KE, Abaryan Z, Laltoo E, Hernandez LM, Gandal MJ, McCracken JT, Thompson PM (2023): White matter microstructure shows sex differences in late childhood: Evidence from 6797 children. Hum Brain Mapp 44: 535–548.

40. Raznahan A, Shaw P, Lalonde F, Stockman M, Wallace GL, Greenstein D, et al. (2011, May 11): How does your cortex grow? The Journal of Neuroscience: The Official Journal of the Society for Neuroscience, vol. 31. pp 7174–7177.

41. Chan JC, Nugent BM, Bale TL (2018): Parental Advisory: Maternal and Paternal Stress Can Impact Offspring Neurodevelopment. Biol Psychiatry 83: 886–894.

42. Brew BK, Lundholm C, Caffrey Osvald E, Chambers G, Öberg S, Fang F, Almqvist C (2022): Early-life adversity due to bereavement and inflammatory diseases in the next generation: A population study in transgenerational stress exposure. Am J Epidemiol 191: 38–48.

43. Yehuda R, Daskalakis NP, Lehrner A, Desarnaud F, Bader HN, Makotkine I, et al. (2014): Influences of maternal and paternal PTSD on epigenetic regulation of the glucocorticoid receptor gene in Holocaust survivor offspring. Am J Psychiatry 171: 872–880.

44. Uematsu A, Matsui M, Tanaka C, Takahashi T, Noguchi K, Suzuki M, Nishijo H (2012): Developmental trajectories of amygdala and hippocampus from infancy to early adulthood in healthy individuals. PLoS One 7: e46970.

45. Dima D, Modabbernia A, Papachristou E, Doucet GE, Agartz I, Aghajani M, et al. (2022): Subcortical volumes across the lifespan: Data from 18,605 healthy individuals aged 3-90 years. Hum Brain Mapp 43: 452–469.

46. White T, Su S, Schmidt M, Kao C-Y, Sapiro G (2010): The development of gyrification in childhood and adolescence. Brain Cogn 72: 36–45.

47. Armstrong E, Schleicher A, Omran H, Curtis M, Zilles K (1995): The ontogeny of human gyrification. Cereb Cortex 5: 56–63.

48. Quezada S, Castillo-Melendez M, Walker DW, Tolcos M (2018): Development of the cerebral cortex and the effect of the intrauterine environment: Cortical development and intrauterine environment. J Physiol 596: 5665–5674.

49. Sasabayashi D, Takahashi T, Takayanagi Y, Suzuki M (2021): Anomalous brain gyrification patterns in major psychiatric disorders: a systematic review and transdiagnostic integration. Transl Psychiatry 11: 176.

50. Mihailov A, Pron A, Lefèvre J, Deruelle C, Desnous B, Bretelle F, et al. (2025): Burst of gyrification in the human brain after birth. Commun Biol 8: 805.

51. Mareckova K, Miles A, Andryskova L, Brazdil M, Nikolova YS (2020): Temporally and sex-specific effects of maternal perinatal stress on offspring cortical gyrification and mood in young adulthood. Hum Brain Mapp 41: 4866–4875.

52. Papini C, Palaniyappan L, Kroll J, Froudist-Walsh S, Murray RM, Nosarti C (2020): Altered Cortical Gyrification in Adults Who Were Born Very Preterm and Its Associations With Cognition and Mental Health. Biol Psychiatry Cogn Neurosci Neuroimaging 5: 640–650.

53. Kühn S, Witt C, Banaschewski T, Barbot A, Barker GJ, Büchel C, et al. (2016): From mother to child: orbitofrontal cortex gyrification and changes of drinking behaviour during adolescence. Addict Biol 21: 700–708.

54. Haukvik UK, Schaer M, Nesvåg R, McNeil T, Hartberg CB, Jönsson EG, et al. (2012): Cortical folding in Broca’s area relates to obstetric complications in schizophrenia patients and healthy controls. Psychol Med 42: 1329–1337.

55. Lu Y-C, Andescavage N, Wu Y, Kapse K, Andersen NR, Quistorff J, et al. (2022): Maternal psychological distress during the COVID-19 pandemic and structural changes of the human fetal brain. Communications Medicine 2: 1–13.

56. Plant DT, Pawlby S, Pariante CM, Jones FW (2018): When one childhood meets another - maternal childhood trauma and offspring child psychopathology: A systematic review. Clin Child Psychol Psychiatry 23: 483–500.

57. Matsudaira I, Yamaguchi R, Taki Y (2023): Transmit Radiant Individuality to Offspring (TRIO) study: investigating intergenerational transmission effects on brain development. Front Psychiatry 14: 1150973.

58. Bernstein DP, Stein JA, Newcomb MD, Walker E, Pogge D, Ahluvalia T, et al. (2003): Development and validation of a brief screening version of the Childhood Trauma Questionnaire. Child Abuse Negl 27: 169–190.

59. Nakajima M, Hori H, Itoh M, Lin M, Kawanishi H, Narita M, Kim Y (2022): Validation of childhood trauma questionnaire-short form in Japanese clinical and nonclinical adults. Psychiatry Res Commun 2: 100065.

60. Assink M, Spruit A, Schuts M, Lindauer R, van der Put CE, Stams G-JJM (2018): The intergenerational transmission of child maltreatment: A three-level meta-analysis. Child Abuse Negl 84: 131–145.

61. Madigan S, Cyr C, Eirich R, Fearon RMP, Ly A, Rash C, et al. (2019): Testing the cycle of maltreatment hypothesis: Meta-analytic evidence of the intergenerational transmission of child maltreatment. Dev Psychopathol 31: 23–51.

62. Kessler RC, Andrews G, Colpe LJ, Hiripi E, Mroczek DK, Normand SLT, et al. (2002): Short screening scales to monitor population prevalences and trends in non-specific psychological distress. Psychol Med 32: 959–976.

63. Furukawa TA, Kawakami N, Saitoh M, Ono Y, Nakane Y, Nakamura Y, et al. (2008): The performance of the Japanese version of the K6 and K10 in the World Mental Health Survey Japan. Int J Methods Psychiatr Res 17: 152–158.

64. Fraley RC, Heffernan ME, Vicary AM, Brumbaugh CC (2011): The Experiences in Close Relationships- Relationship Structures questionnaire: a method for assessing attachment orientations across relationships. Psychol Assess 23: 615–625.

65. Komura K, Murakami T, Toda K (2016): Validation of a Japanese version of the Experience in Close Relationship- Relationship Structure. Shinrigaku Kenkyu 87: 303–313.

66. Matsudaira I, Yamaguchi R, Taki Y (2025): Parent-offspring brain similarity: Specificities and commonalities among sex combinations-the transmit radiant individuality to offspring study. iScience 28: 112936.

67. Glasser MF, Sotiropoulos SN, Wilson JA, Coalson TS, Fischl B, Andersson JL, et al. (2013): The minimal preprocessing pipelines for the Human Connectome Project. Neuroimage 80: 105–124.

68. Tustison NJ, Avants BB, Cook PA, Zheng Y, Egan A, Yushkevich PA, Gee JC (2010): N4ITK: improved N3 bias correction. IEEE Trans Med Imaging 29: 1310–1320.

69. Isensee F, Schell M, Pflueger I, Brugnara G, Bonekamp D, Neuberger U, et al. (2019): Automated brain extraction of multisequence MRI using artificial neural networks. Hum Brain Mapp 40: 4952–4964.

70. Fischl B (2012): FreeSurfer. Neuroimage 62: 774–781.

71. Schaer M, Cuadra MB, Tamarit L, Lazeyras F, Eliez S, Thiran J-P (2008): A surface-based approach to quantify local cortical gyrification. IEEE Trans Med Imaging 27: 161–170.

72. Glasser MF, Coalson TS, Robinson EC, Hacker CD, Harwell J, Yacoub E, et al. (2016): A multi-modal parcellation of human cerebral cortex. Nature 536: 171–178.

73. Benjamini Y, Hochberg Y (1995): Controlling the false discovery rate: A practical and powerful approach to multiple testing. J R Stat Soc Series B Stat Methodol 57: 289–300.

74. Imai K, Keele L, Tingley D (2010): A general approach to causal mediation analysis. Psychol Methods. 10.1037/a0020761

75. Tingley D, Yamamoto T, Hirose K, Keele L, Imai K (2014): Mediation:RPackage for causal mediation analysis. J Stat Softw 59: 1–38.

76. R Core Team (2023): R: A Language and Environment for Statistical Computing. Vienna, Austria: R Foundation for Statistical Computing. Retrieved from https://www.R-project.org/

77. Mowinckel AM, Vidal-Piñeiro D (2020): Visualization of Brain Statistics With R Packages ggseg and ggseg3d. Advances in Methods and Practices in Psychological Science 3: 466–483.

78. Mychasiuk R, Gibb R, Kolb B (2012): Prenatal stress alters dendritic morphology and synaptic connectivity in the prefrontal cortex and hippocampus of developing offspring. Synapse 66: 308– 314.

79. Zhang Z, Li N, Chen R, Lee T, Gao Y, Yuan Z, et al. (2021): Prenatal stress leads to deficits in brain development, mood related behaviors and gut microbiota in offspring. Neurobiol Stress 15: 100333.

80. Marchisella F, Creutzberg KC, Begni V, Sanson A, Wearick-Silva LE, Tractenberg SG, et al. (2021): Exposure to prenatal stress is associated with an excitatory/inhibitory imbalance in rat prefrontal cortex and amygdala and an increased risk for emotional dysregulation. Front Cell Dev Biol 9: 653384.

81. Yun HJ, Lee HJ, Lee JY, Tarui T, Rollins CK, Ortinau CM, et al. (2022): Quantification of sulcal emergence timing and its variability in early fetal life: Hemispheric asymmetry and sex difference. Neuroimage 263: 119629.

82. Wade M, Wright L, Finegold KE (2022): The effects of early life adversity on children’s mental health and cognitive functioning. Transl Psychiatry 12: 1–12.

83. Teicher MH, Samson JA, Anderson CM, Ohashi K (2016): The effects of childhood maltreatment on brain structure, function and connectivity. Nat Rev Neurosci 17: 652–666.

84. McLaughlin KA, Weissman D, Bitrán D (2019): Childhood adversity and neural development: A systematic review. Annu Rev Dev Psychol 1: 277–312.

85. Kim MJ, Loucks RA, Palmer AL, Brown AC, Solomon KM, Marchante AN, Whalen PJ (2011): The structural and functional connectivity of the amygdala: from normal emotion to pathological anxiety. Behav Brain Res 223: 403–410.

86. Morawetz C, Basten U (2024): Neural underpinnings of individual differences in emotion regulation: A systematic review. Neurosci Biobehav Rev 162: 105727.

87. Wilcox CE, Pommy JM, Adinoff B (2016): Neural circuitry of impaired emotion regulation in substance use disorders. Am J Psychiatry 173: 344–361.

88. Gallucci J, Secara MT, Chen O, Oliver LD, Jones BDM, Marawi T, et al. (2024): A systematic review of structural and functional magnetic resonance imaging studies on the neurobiology of depressive symptoms in schizophrenia spectrum disorders. Schizophrenia (Heidelb) 10: 59.

89. Kolesar TA, Bilevicius E, Wilson AD, Kornelsen J (2019): Systematic review and meta-analyses of neural structural and functional differences in generalized anxiety disorder and healthy controls using magnetic resonance imaging. NeuroImage Clin 24: 102016.

90. Guma E, Chakravarty MM (2025): Immune alterations in the intrauterine environment shape offspring brain development in a sex-specific manner. Biol Psychiatry 97: 12–27.

91. Mueller BR, Bale TL (2008): Sex-specific programming of offspring emotionality after stress early in pregnancy. J Neurosci 28: 9055–9065.

92. Howerton CL, Morgan CP, Fischer DB, Bale TL (2013): O-GlcNAc transferase (OGT) as a placental biomarker of maternal stress and reprogramming of CNS gene transcription in development. Proc Natl Acad Sci U S A 110: 5169–5174.

93. Howerton CL, Bale TL (2014): Targeted placental deletion of OGT recapitulates the prenatal stress phenotype including hypothalamic mitochondrial dysfunction. Proc Natl Acad Sci U S A 111: 9639– 9644.

94. Clifton VL (2010): Review: Sex and the Human Placenta: Mediating Differential Strategies of Fetal Growth and Survival. Placenta 31: S33–S39.

95. Bosquet Enlow M, Sideridis G, Bollati V, Hoxha M, Hacker MR, Wright RJ (2019): Maternal cortisol output in pregnancy and newborn telomere length: Evidence for sex-specific effects. Psychoneuroendocrinology 102: 225–235.

96. de Tychey C, Briançon S, Lighezzolo J, Spitz E, Kabuth B, de Luigi V, et al. (2008): Quality of life, postnatal depression and baby gender. J Clin Nurs 17: 312–322.

97. Myers S, Johns SE (2019): Male infants and birth complications are associated with increased incidence of postnatal depression. Soc Sci Med 220: 56–64.

98. Cowell W, Colicino E, Askowitz T, Nentin F, Wright RJ (2021): Fetal sex and maternal postpartum depressive symptoms: findings from two prospective pregnancy cohorts. Biol Sex Differ 12: 6.

99. Troisi R, Potischman N, Roberts J, Siiteri P, Daftary A, Sims C, Hoover RN (2003): Associations of maternal and umbilical cord hormone concentrations with maternal, gestational and neonatal factors (United States). Cancer Causes Control 14: 347–355.

100. Toriola AT, Vääräsmäki M, Lehtinen M, Zeleniuch-Jacquotte A, Lundin E, Rodgers K-G, et al. (2011): Determinants of maternal sex steroids during the first half of pregnancy. Obstet Gynecol 118: 1029– 1036.

101. Letourneau N, Dewey D, Kaplan BJ, Ntanda H, Novick J, Thomas JC, et al. (2019): Intergenerational transmission of adverse childhood experiences via maternal depression and anxiety and moderation by child sex. J Dev Orig Health Dis 10: 88–99.

102. Jawaid A, Kunzi M, Mansoor M, Khan ZY, Abid A, Taha M, et al. (2020, August 14): Distinct microRNA signature in human serum and germline after childhood trauma. MedRxiv. medRxiv, p 2020.08.11.20168393.

103. Merrill SM, Moore SR, Gladish N, Giesbrecht GF, Dewey D, Konwar C, et al. (2021): Paternal adverse childhood experiences: Associations with infant DNA methylation. Dev Psychobiol 63: e22174.

104. Stoltenborgh M, Bakermans-Kranenburg MJ, Alink LRA, van IJzendoorn MH (2015): The prevalence of child maltreatment across the globe: Review of a series of meta-analyses: Prevalence of child maltreatment across the globe. Child Abuse Rev 24: 37–50.

105. Everaerd D, Klumpers F, Zwiers M, Guadalupe T, Franke B, van Oostrom I, et al. (2016): Childhood abuse and deprivation are associated with distinct sex-dependent differences in brain morphology. Neuropsychopharmacology 41: 1716–1723.

106. Teicher MH, Anderson CM, Ohashi K, Khan A, McGreenery CE, Bolger EA, et al. (2018): Differential effects of childhood neglect and abuse during sensitive exposure periods on male and female hippocampus. Neuroimage 169: 443–452.

107. Prachason T, Mutlu I, Fusar-Poli L, Menne-Lothmann C, Decoster J, van Winkel R, et al. (2024): Gender differences in the associations between childhood adversity and psychopathology in the general population. Soc Psychiatry Psychiatr Epidemiol 59: 847–858.

108. Zhu S, Cheng S, Liu W, Ma J, Sun W, Xiao W, et al. (2025): Gender differences in the associations of adverse childhood experiences with depression and anxiety: A systematic review and meta-analysis. J Affect Disord 378: 47–57.

109. Kawata NYS, Fujisawa TX, Makita K, Yao A, Okazawa H, Tomoda A (2025): White matter microstructure abnormalities in children experiencing neglect without other forms of maltreatment. Sci Rep 15: 27282.

110. Franklin TB, Russig H, Weiss IC, Gräff J, Linder N, Michalon A, et al. (2010): Epigenetic transmission of the impact of early stress across generations. Biol Psychiatry 68: 408–415.

111. Saavedra-Rodríguez L, Feig LA (2013): Chronic social instability induces anxiety and defective social interactions across generations. Biol Psychiatry 73: 44–53.

112. He N, Kong Q-Q, Wang J-Z, Ning S-F, Miao Y-L, Yuan H-J, et al. (2016): Parental life events cause behavioral difference among offspring: Adult pre-gestational restraint stress reduces anxiety across generations. Sci Rep 6: 39497.

113. Mashoodh R, Habrylo IB, Gudsnuk K, Champagne FA (2023): Sex-specific effects of chronic paternal stress on offspring development are partially mediated via mothers. Horm Behav 152: 105357.

114. Masson BA, Kiridena P, Lu D, Kleeman EA, Reisinger SN, Qin W, et al. (2025): Depletion of the paternal gut microbiome alters sperm small RNAs and impacts offspring physiology and behavior in mice. Brain Behav Immun 123: 290–305.

115. Gabory A, Attig L, Junien C (2009): Sexual dimorphism in environmental epigenetic programming. Mol Cell Endocrinol 304: 8–18.

116. Gabory A, Roseboom TJ, Moore T, Moore LG, Junien C (2013): Placental contribution to the origins of sexual dimorphism in health and diseases: sex chromosomes and epigenetics. Biol Sex Differ 4: 5.

117. Rosenfeld CS (2015): Sex-specific placental responses in fetal development. Endocrinology 156: 3422–3434.

118. Buss C, Entringer S, Moog NK, Toepfer P, Fair DA, Simhan HN, et al. (2017): Intergenerational Transmission of Maternal Childhood Maltreatment Exposure: Implications for Fetal Brain Development. J Am Acad Child Adolesc Psychiatry 56: 373–382.

119. Jones DK, Christiansen KF, Chapman RJ, Aggleton JP (2013): Distinct subdivisions of the cingulum bundle revealed by diffusion MRI fibre tracking: implications for neuropsychological investigations. Neuropsychologia 51: 67–78.

120. Bubb EJ, Metzler-Baddeley C, Aggleton JP (2018): The cingulum bundle: Anatomy, function, and dysfunction. Neurosci Biobehav Rev 92: 104–127.

121. Ahmed SP, Bittencourt-Hewitt A, Sebastian CL (2015): Neurocognitive bases of emotion regulation development in adolescence. Dev Cogn Neurosci 15: 11–25.

122. Marusak HA, Thomason ME, Peters C, Zundel C, Elrahal F, Rabinak CA (2016): You say “prefrontal cortex” and I say “anterior cingulate”: meta-analysis of spatial overlap in amygdala-to-prefrontal connectivity and internalizing symptomology. Transl Psychiatry 6: e944.

123. Schmaal L, Hibar DP, Sämann PG, Hall GB, Baune BT, Jahanshad N, et al. (2017): Cortical abnormalities in adults and adolescents with major depression based on brain scans from 20 cohorts worldwide in the ENIGMA Major Depressive Disorder Working Group. Mol Psychiatry 22: 900–909.

124. Hiser J, Koenigs M (2018): The multifaceted role of the ventromedial prefrontal cortex in emotion, decision making, social cognition, and psychopathology. Biol Psychiatry 83: 638–647.

125. Li Q, Zhao Y, Chen Z, Long J, Dai J, Huang X, et al. (2020): Meta-analysis of cortical thickness abnormalities in medication-free patients with major depressive disorder. Neuropsychopharmacology 45: 703–712.

126. Rowold ED, Schulze L, Van der Auwera S, Grabe HJ (2017): Paternal transmission of early life traumatization through epigenetics: Do fathers play a role? Med Hypotheses 109: 59–64.

127. Godbout N, Daspe M-È, Runtz M, Cyr G, Briere J (2019): Childhood maltreatment, attachment, and borderline personality-related symptoms: Gender-specific structural equation models. Psychol Trauma 11: 90–98.

128. Folger AT, Eismann EA, Stephenson NB, Shapiro RA, Macaluso M, Brownrigg ME, Gillespie RJ (2018): Parental adverse childhood experiences and offspring development at 2 years of age. Pediatrics 141. 10.1542/peds.2017-2826

129. Pietikäinen JT, Kiviruusu O, Kylliäinen A, Pölkki P, Saarenpää-Heikkilä O, Paunio T, Paavonen EJ (2020): Maternal and paternal depressive symptoms and children’s emotional problems at the age of 2 and 5 years: a longitudinal study. J Child Psychol Psychiatry 61: 195–204.

130. Zhen-Duan J, Cruz-Gonzalez M, Diaz J, Sánchez M, Park I, Alvarez K, et al. (2025): Intergenerational continuity of adverse childhood experiences among Mexican-origin families: Examination of intra and extra-familial adversities. Fam Process 64: e13091.

131. Li R, Jia L, Zha J, Wang X, Huang Y, Tao X, Wan Y (2025): Association of maternal and paternal adverse childhood experiences with emotional and behavioral problems among preschool children. Eur Child Adolesc Psychiatry 34: 1111–1123.

132. Gehred MZ, Knodt AR, Ambler A, Bourassa KJ, Danese A, Elliott ML, et al. (2021): Long-term neural embedding of childhood adversity in a population-representative birth cohort followed for 5 decades. Biol Psychiatry 90: 182–193.

